# A population of descending neurons that regulate the flight motor of Drosophila

**DOI:** 10.1101/2021.08.05.455281

**Authors:** Shigehiro Namiki, Ivo G. Ros, Carmen Morrow, William J. Rowell, Gwyneth M. Card, Wyatt Korff, Michael H. Dickinson

## Abstract

Like many insect species, *Drosophila melanogaster* are capable of maintaining a stable flight trajectory for periods lasting up to several hours(1, 2). Because aerodynamic torque is roughly proportional to the fifth power of wing length(3), even small asymmetries in wing size require the maintenance of subtle bilateral differences in flapping motion to maintain a stable path. Flies can even fly straight after losing half of a wing, a feat they accomplish via very large, sustained kinematic changes to the both damaged and intact wings(4). Thus, the neural network responsible for stable flight must be capable of sustaining fine-scaled control over wing motion across a large dynamic range. In this paper, we describe an unusual type of descending neurons (DNg02) that project directly from visual output regions of the brain to the dorsal flight neuropil of the ventral nerve cord. Unlike most descending neurons, which exist as single bilateral pairs with unique morphology, there is a population of at least 15 DNg02 cell pairs with nearly identical shape. By optogenetically activating different numbers of DNg02 cells, we demonstrate that these neurons regulate wingbeat amplitude over a wide dynamic range via a population code. Using 2-photon functional imaging, we show that DNg02 cells are responsive to visual motion during flight in a manner that would make them well suited to continuously regulate bilateral changes in wing kinematics. Collectively, we have identified a critical set of DNs that provide the sensitivity and dynamic range required for flight control.

## Results

Within a fly’s nervous system, sensory information from the brain is conveyed to motor regions of the ventral nerve cord (VNC) by several hundred pairs of descending neurons (DNs) that are roughly stratified into a dorsal pathway that projects to flight motor neuropils and a ventral pathway that project to leg neuromeres(5, 6) (Figure 1A, B). Whereas most of the DNs are single pairs of bilateral cells with unique morphology, a small number of DNs constitute larger sets of homomorphic neurons. To identify classes of DNs that might be involved in flight control, we conducted an activation screen in which we expressed CsChrimson(7) in one GAL4 line and 48 split-GAL4 lines(8) that collectively target 29 different dorsally projecting DNs that innervate the wing and haltere neuropils and tectulum of the VNC(6), along with one control split-GAL4 line that does not drive expression in any neuron (“empty”). In each trial, we aligned tethered, flying flies within a machine vision system to measure wingbeat amplitude(9). To promote stable flight during each trial, we presented the flies with a pattern of vertical stripes presented on an LED array(10) that covered 212° of their frontal field of view. While the flies regulated the angular velocity of the visual pattern under closed-loop conditions, we presented a brief, 100 ms pulse of 617 nm light to activate the targeted DNs (Fig. 1C, D). These 50 lines varied not only with respect to the cells they labeled, but also the sparsity of expression. Nevertheless, a clear pattern emerged when comparing the results across lines (Fig. 1E). Of the 13 lines in which CsChrimson activation resulted in the largest change in wingbeat amplitude, 11 targeted members of the same class of neurons, DNg02.

**Figure 1.**
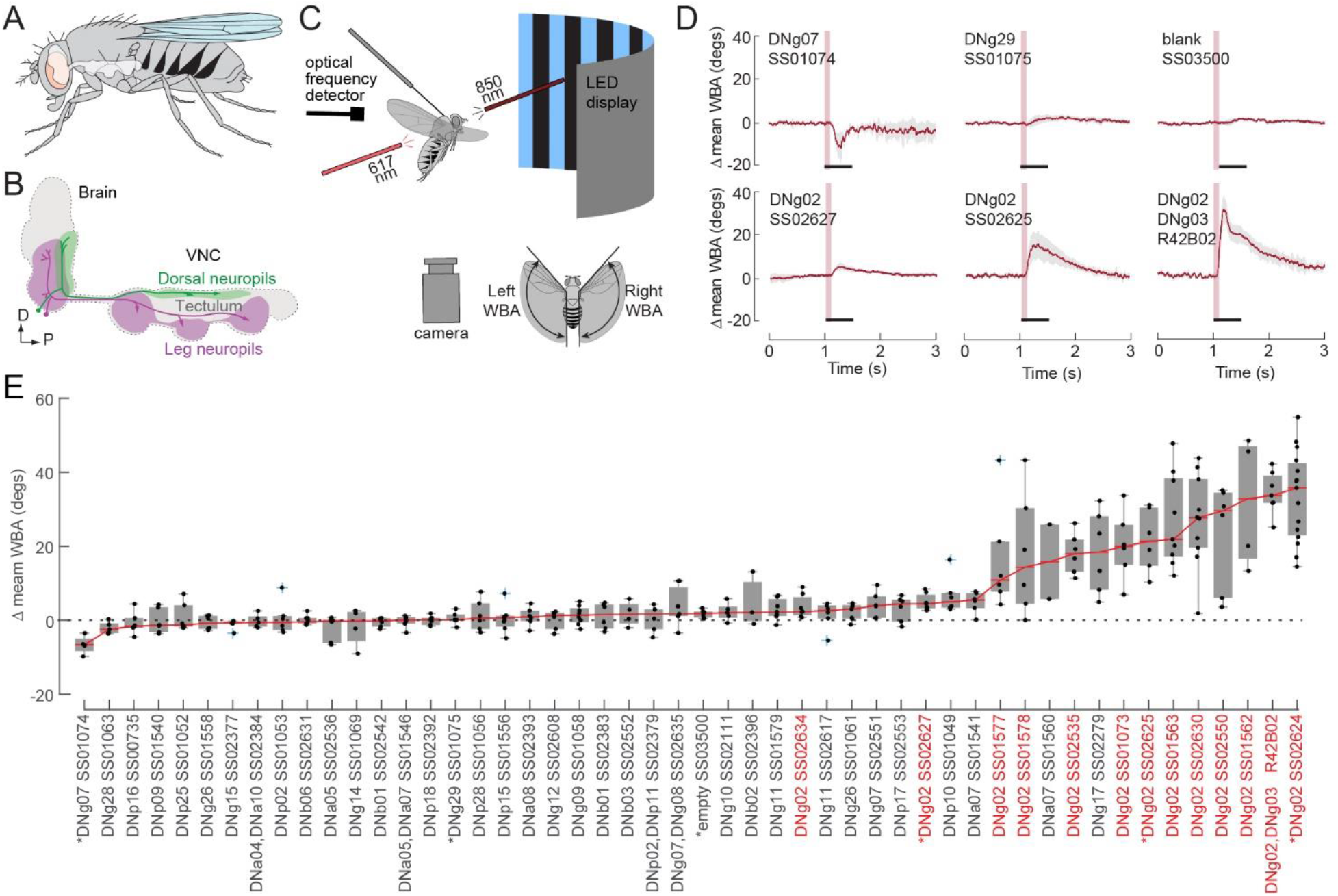
Experimental methods for optogenetic activation and screen results. (A) Cartoon showing location of central nervous system in a standing fly. (B) Descending neurons (DNs) are stratified into two main groups: a ventral group (magenta) that innervates the three leg neuromeres and a dorsal group (green) that innervates the dorsal neuropils associated with the neck, wings, and halteres. (C) Cartoon showing experimental set-up (not drawn to scale). A fiber optic cable delivering 617 nm light is positioned behind a tethered fly aimed at the thorax. Two fiber optic cables (only one is shown) deliver continuous near-IR light (850 nm) to illuminate the left and right stroke planes of the fly. The fly is centered within a curved visual display of blue (470 nm) LEDs upon which a striped drum (spatial frequency = 36o) was presented under closed-loop conditions. An image of the fly is captured with an upward facing camera and analyzed using a real-time machine vision system that measures the angular extent of the left and right wingbeat amplitudes. In all experiments, we assumed that the rearward reversal angle was parallel to the body axis and remained constant. An optical detector that recorded fluctuations in IR light was used to record wingbeat frequency. (D) Results of 100 ms pulses of CsChrimson activation on the change in the mean of the left and right wingbeat amplitude of the wings in 6 of the 50 lines tested; red line and gray area indicate the mean response and the standard deviation envelope, respectively. One line (SS01074), which targets the DNg07 neuron, was distinct in that it elicited a consistent decrease in wingbeat amplitude. Most targeted cells, such as the DNg29 neuron shown in Figure 1 (labeled by SS01075) did not elicit a detectable change in wingbeat amplitude. Control flies in which the UAS-CsChrimson line was crossed with an empty vector split-GAL4 line (blank, SS03500) exhibited no response to the 617 nm light. Bottom row: Example traces of different driver lines that target the DNg02 cells. For each trial, we determined the average wingbeat amplitude (WBA) of the left and right wings over the 0.5 second period prior to stimulus onset and subtracted this value from the entire trace to create a zero baseline. The response to optogenetic activation was calculated as the average value of WBA over the 0.5 second period starting with stimulus onset. (E) Overview of the entire screen of 50 lines, ranked from left to right according to the magnitude of the optogenetic effect on wingbeat amplitude. Each trial was scored by averaging the change in the mean of wingbeat amplitude over the 500 ms period beginning with the onset of the light pulse (indicated by black bars in B). For each line, the median response is indicated by a red line. The median value from each individual fly is indicated by a black dot, the box and whisker plots indicate the interquartile range and extreme values for the flies tested; outliers are indicated by blue crosses. Lines that target DNg02 neurons are indicated by red font. Asterisks indicate lines for which the responses are plotted in panel D.

The DNg02s were previously identified anatomically(6) and consist of a cluster of at least 15 cell pairs with nearly identical morphology, the largest of the population-type class of DNs identified so far. Their small, spindly cell bodies reside in a cluster at the ventral edge of the gnathal ganglion (GNG; Figure 2A). The primary neurites of the DNg02s run ventrally along the edge of the GNG before taking a hairpin turn and ascending dorsally, where each cell arborizes in a hemi-circle around the esophageal foramen. The cells’ terminals reside within a set of five contiguous neuropils consisting of the inferior bridge (IB), inferior clamp (ICL), superior posterior slopes (SPS), inferior posterior slope (IPS), as well as the gnathal ganglion (GNG)(11). Synaptotagmin labeling and fine morphology suggest that processes within the IB and GNG are outputs, whereas those within the IC, SPS, and IPS are inputs(6).

**Figure 2.**
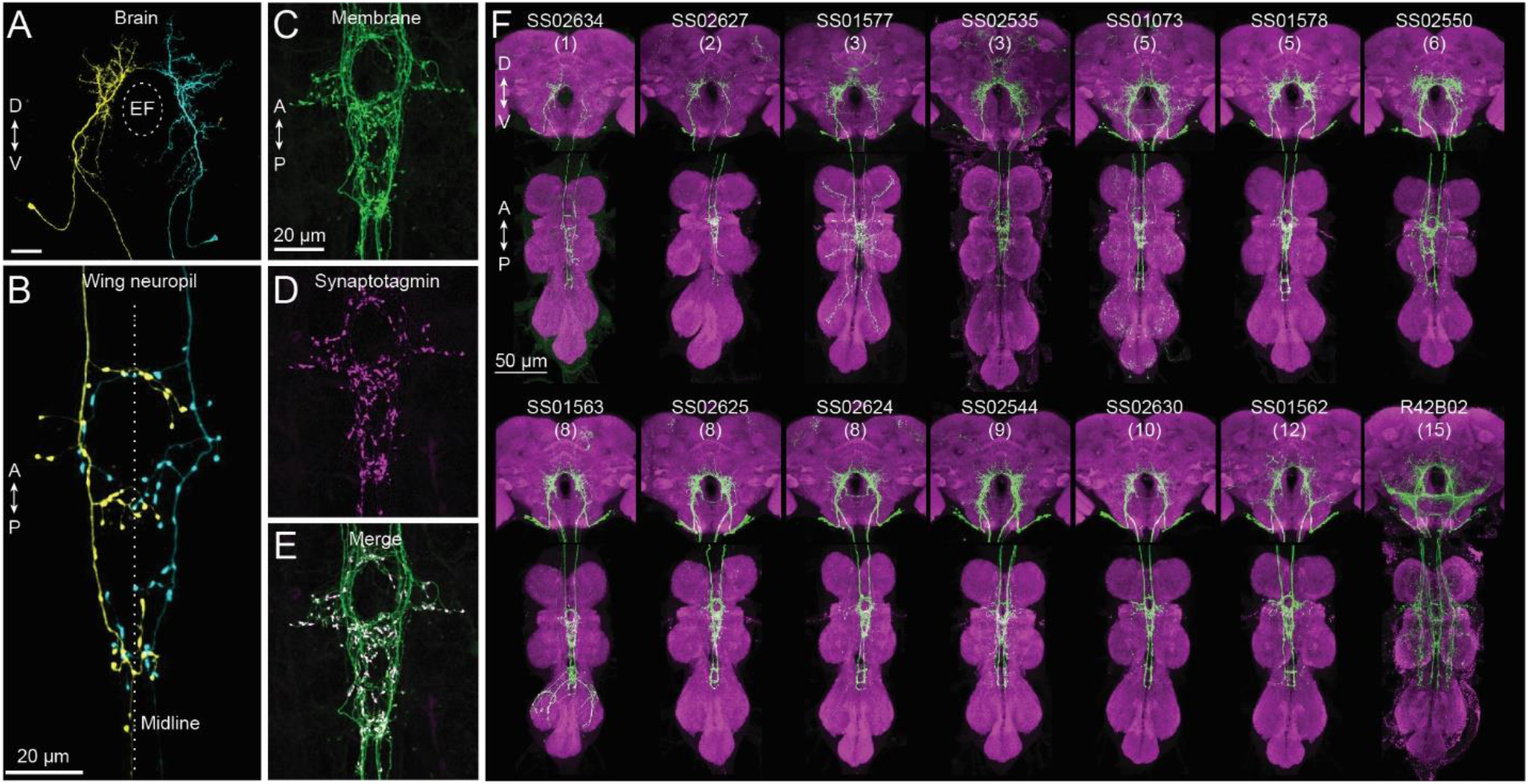
Morphology of DNg02 cells and split-GAL4 driver lines targeting different numbers of neurons. (A) MCFO of SS02627 driver line showing one left and one right DNg02 neuron in the brain; approximate position of esophageal foramen (EF) is indicated by dotted ellipse. (B) Projections of the same neurons in the wing neuropil. The neurons possess punctate terminals on both sides of the VNC midline. (C) Enlarged view of wing neuropil region of SS01578 showing GFP expression in green. (D) Same region as in B, showing synaptotagmin staining in magenta. (E) Merger of images from B and C; DNg02 projections in the wing neuropil contain many output terminals. (F) Expression pattern in 14 different driver lines that target DNg02 neurons. Membrane-targeted GFP expression is shown in green, nc82 staining is shown in magenta. The number of neurons targeted in each line is shown in parenthesis below the line name.

After descending ipsilaterally down the neck connectives, the DNg02 cells exhibit a distinct pattern of arborization in the dorsal VNC (Figure 2B). Collectively, the population of cells forms a compact ‘figure-of-eight’ shape within the dorsal flight neuropil – a pattern also repeated in the haltere neuropil (Figure 2F). Visualization of the morphology of individual neurons using the multi-color flip out (MCFO) expression system(12) indicates that the arborizations of DNg02s remain restricted to the ipsilateral side within the brain, whereas the terminals in the VNC cross the midsagittal plane (Figure 2A,B). Large synaptotagmin-positive boutons are distributed diffusely throughout the projections in the wing and haltere neuropils (Figure 2C-E), consistent with output synapses in these regions.

The large number of DNg02 cells suggests that they might underlie some unique and critical function within the flight motor system. One hypothesis is that the DNg02s act on wing motion via a population code(13, 14), such that the precise kinematic output of the wings depends in part on the number of DNg02 cells that are active as well as the level of activity within individual cells. To test this hypothesis, we used 13 separate split-GAL4 lines that targeted different subsets of DNg02 cells with little or no expression in off-target neurons (Figure 2F). In addition, we evaluated one GAL4 line (GMR42B02) that targets 15 DNg02 cell pairs. As a control, we tested the empty split-GAL4 line (SS03500). Because the number of DNg02 cell pairs labeled in these 14 lines varied from 0 to 15, we were able to test the influence of population activity by driving each line independently and comparing the magnitude of the effect on wingbeat amplitude during flight. As shown in Figure 3A, we found a strongly linear relationship (r^2^ = 0.7394) between the number of DNg02 cells present in each line and the magnitude of the change in wingbeat amplitude elicited by activation.

**Figure 3.**
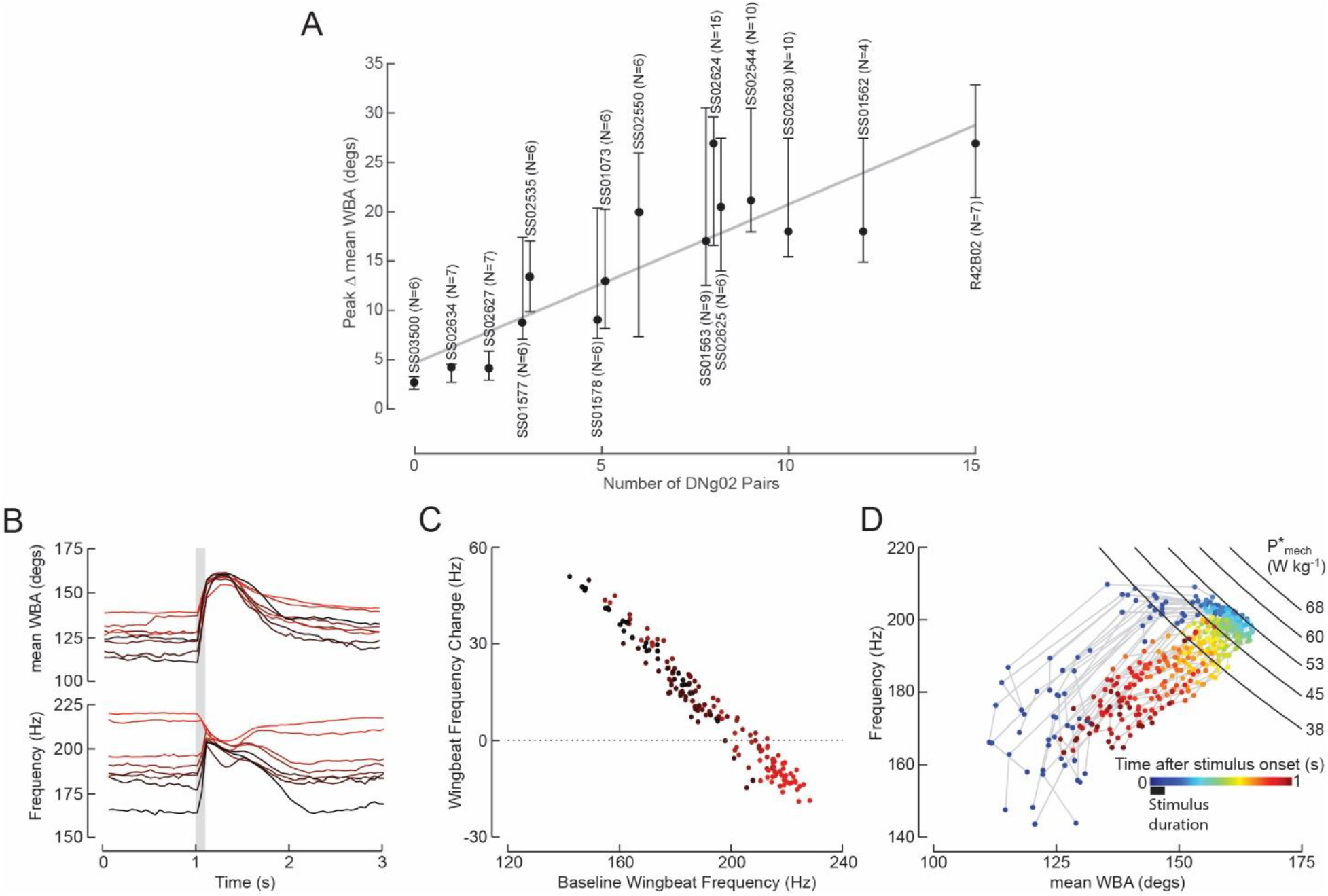
Wingbeat amplitude appears to be regulated via a population code of DNg02 neurons. (A) Peak change in the mean of the left and right WBA elicited by CsChrimson activation in 15 driver lines that target different numbers of DNg02 neurons; black circle indicates median value for each line, vertical bars indicate interquartile range. The identity of each line is labeled by vertical text above or below the data. Three sets of lines are plotted close together because they target the same number of neurons (SS01577, SS02535: 3 cells; SS01578, SS01073: 5 cells; SS01563, SS02624, SS02625: 8 cells). The number of cell pairs activated correlates positively with the magnitude of the changes in mean WBA (r2 = 0.7394, based on median values). (B) Time series traces for changes in mean ((left+right)/2) wingbeat amplitude (top) and frequency (bottom) elicited by optogenetic activation of DNg02 cells in the SS02625 driver line. Each trace represents the mean response for one fly. Each fly (N=8) is indicated by color, with the variation from red to black scaled according to the level of background wingbeat frequency. Note that whereas the background, pre-stimulus wingbeat angle varied among flies, the peak level elicited by CsChrimson activation was quite similar among flies. Wingbeat frequency also showed a large variation across flies before and after the stimulus, and a decreased variation during activation. Note that wingbeat frequency tends to fall during optogenetic activation following an initial rise. (C) Change in wingbeat frequency during optogenetic activation plotted against the pre-stimulus baseline; color scheme indicates identity of flies plotted in B. Note the inverse relationship; when the background level of wingbeat frequency is above ~205 Hz, optogenetic activation elicits a drop in frequency below baseline. (H) Wingbeat frequency plotted against mean ((left+right)/2) wingbeat amplitude for every activation trial of an example fly from panels B and C. For each trial, the time after stimulus onset is encoded by color; the duration of the stimulus is indicated by the black bar below the color scale. At stimulus onset, both wingbeat amplitude and frequency rise; however, after WBA reaches a value of ~160o, further increases in wingbeat amplitude are accompanied by a decrease in wingbeat frequency. The black curves show isolines for muscle mass specific mechanical power (P*mech) in the frequency-amplitude plane; see text for details.

We further explored the effects of optogenetic activation on a subset of driver lines, focusing on SS02625, a line that targets 8 DNg02 cells (Figure 2F). Whereas individual flies varied with respect to their background level of wingbeat amplitude prior to optogenetic activation (Figure 3B, top traces), the peak level of wingbeat amplitude typically reached the same approximate value for each fly during activation. This result suggests that the level of activation we applied elicited a saturating excitation of the DNg02 neurons within this line. In such cases, we also observed an intriguing pattern of changes in wingbeat frequency elicited by the excitation (Figure 3B, bottom traces). In most cases, DNg02 activation resulted in a correlated rise in both wingbeat amplitude and frequency, but in some instances the increase in amplitude was accompanied by a net decrease in frequency. Whether activation elicited a decrease or increase in wingbeat frequency was not random, but rather depended on the level of wingbeat frequency prior to activation. In particular, individuals that flew with a lower wingbeat frequency exhibited a large increase in frequency upon activation, whereas those that flew with a higher wingbeat frequency exhibited a decrease (Figure 3C). A likely explanation for this peculiar trend emerges from plotting instantaneous wingbeat frequency against wingbeat amplitude throughout the time course of optogenetic activation for all the trials of an individual fly (Figure 3C). In each trial, CsChrimson activation evoked an initial rapid rise in both wingbeat amplitude and frequency; however, once the mean wingbeat amplitude of the two wings reached a value of about 160°, further increases in amplitude were accompanied by a small decline in frequency (see also traces in bottom panel of Figure 3B). Thus, when the DNg02 cells within the fly were maximally activated, we observed an inverse relationship between wingbeat frequency and amplitude.

The time-varying relationship between wingbeat amplitude and frequency (Figure 3D) bear a striking resemblance to data presented in a former study on the metabolic power requirements for flight(15), in which changes in wingbeat amplitude and frequency were elicited by upward and downward visual motion rather than optogenetic activation of DNs. That study presented a model in which the mass specific mechanical power (*P*^*^_mech_) delivered by the flight muscles sustains the sum of induced power (the cost of lift) and profile power (the cost of drag) during flight. Profile power, which is the dominant term, is proportional to the product of the wingbeat frequency and amplitude cubed(3). Thus, their model predicts that when the mechanical power generated by the flight muscles is constant—as it is when the asynchronous flight muscles are maximally activated—any increase in wingbeat amplitude must be accompanied by a decrease in frequency and vice versa. To analyze whether DNg02 activation may elicit near maximal power output, we superimposed isolines for mechanical power in the frequency-amplitude plane, using equations from the prior study(15). The precise values of these isolines are only approximate, because they are based on average morphometric data for wing length, wing mass, and body mass of the flies used in the prior study(15); we did not take those measurements on the flies used in our experiments. However, the salient observation is that the set of amplitude-frequency values elicited during peak DNg02 activation are bounded by the shape of the power isolines (e.g. *P*^*^_mech_~ 105 W kg^−1^), which thus enforces the observed inverse relationship between wingbeat frequency and wingbeat amplitude. These data suggest that optogenetic activation of this particular driver line (SS02625) results in the production of peak mechanical power, presumably via activation of the motor neurons of the large indirect flight muscles.

So far, our activation experiments suggest that the population of DNg02 cells might function together to regulate flight power like a throttle, by controlling wingbeat amplitude in a bilateral fashion via symmetric activation of power and steering muscle motor neurons. However, this result may simply reflect the fact that optogenetic activation simultaneously excites both the left and right DNs in each fly. To test if left and right DNg02 cells might operate independently during steering maneuvers in flight, we performed 2-photon functional imaging from the dendritic region of the neurons in one of the DNg02 driver lines (SS02535) in tethered flying flies using GCaMP6f as an activity indicator (Figure 4A). In preparing the flies for recording, we dissected a window in the head capsule just dorsal to the esophageal foramen, which allowed us to image neurons on the left and right side of the brain simultaneously (Figure 4B). Because our goal was to test whether left and right DNg02 cells might be active independently, we subjected the flies to an array of different visual patterns during flight, chosen to elicit both symmetrical and asymmetrical wingbeat responses. These stimulus epochs included a widefield pattern that moved upward or downward, yaw motion to the left or right, a stripe oscillating on the left or right, roll motion to the left or right, an expanding object on the left or right, progressive and regressive motion, and closed loop stripe fixation (Figure 4C). The net results from this array of visual patterns was consistent in that they demonstrated unequivocally that at least some DNg02 cells can respond differently on the left and right sides of the brain. This result was most apparent in yaw stimuli (Figure 4D, left traces), during which rightward motion elicited an increase in activity of the right DNg02 cells and a simultaneous decrease in activity of the left DNg02 cells (and vice versa for leftward motion). The changes in cell fluorescence were accompanied by the expected asymmetric changes in wingbeat amplitude for a yaw response. In contrast, bilaterally symmetrical visual patterns, such as regressive visual motion, elicited synchronous changes in activity of the right and left DNg02 cells, accompanied by symmetrical changes in wingbeat amplitude (Figure 4D, rightmost traces). These results indicate that the DNg02 cells can operate independently on the left and right side of the brain in an asymmetrical or symmetrical fashion, depending on the pattern of visual input. Across all recordings, we measured a strong correlation between the DNg02 cells and the wingbeat amplitude of the contralateral wing, and a weaker anti-correlation with the wingbeat amplitude of the ipsilateral wing (Figure 4E). These patterns were readily apparent in individual recordings, in which we determined the correlation coefficient between changes in fluorescence (ΔF/F) and wingbeat amplitude for each pixel in the fluorescence image during a 20 second flight epoch (Figure 4F). A pixel pattern that corresponds to the arbor of the right DNg02 cells was highly correlated with left wingbeat amplitude, whereas a pixel pattern corresponding to the left DNg02 cells was highly correlated with right wingbeat amplitude.

**Figure 4.**
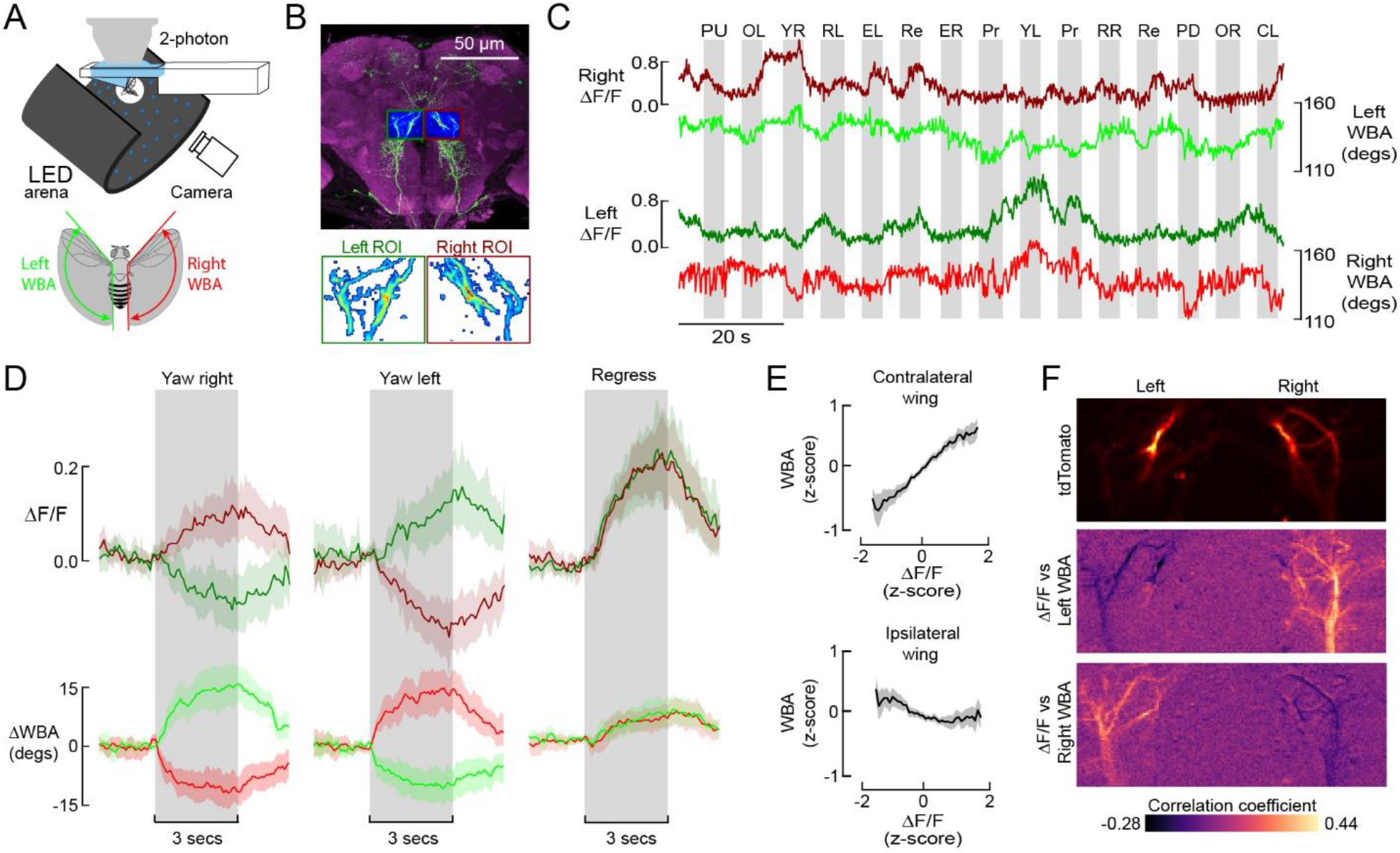
Left and right DNg02 neurons can act independently to regulate wingbeat amplitude. **(A)** Schematic showing fly tethered to a 2-photon microscope surrounded by an LED array for presentation of visual stimuli and a camera for tracking wing motion. Inset: a real-time machine vision system tracks wingbeat amplitude (WBA) the left (green) and right (red) wings of the fly. (**B**) During functional imaging, we captured DNg02 activity within left (dark green) and right (dark red) regions of interest, allowing us to measure simultaneous activity across populations of bilateral cells. The background image is replotted from Figure 2F. Lower inset: within the left and right ROIs, we used a standard deviation threshold to create a mask within which we measured changes in GCaMP6f fluorescence (ΔF/F) while the flying fly was subjected to different patterns of visual motion. (**C**) Example time traces of left and right DNg02 activity (measured as DF/F) along with changes in wingbeat amplitude during presentation of a panel of different visual stimuli (PU = pattern up, OL = stripe oscillating on right, YR = yaw right, RL = roll left, EL = expansion right, Re = Progressive motion, YL = yaw left, Pr = progressive motion, YL = yaw left, RR = roll right, PD = pattern down, OR = stripe oscillating on right, CL = closed loop with stripe). (**D**) Averaged responses to three most informative patterns of optic flow: yaw right, yaw left, and regressive motion (N = 20 flies). Top row shows baseline-subtracted ΔF/F signals from left and right ROIs (dark green and dark red, respectively). Bottom row shows baseline-subtracted wingbeat amplitude signals for left and right wings (green and red, respectively). All data are presented as mean (solid line) and a boot-strapped 95% CI for the mean (shaded area); the 3-second period of stimulus presentation is indicated by the grey patch. (**E**) Wingbeat amplitude of the contralateral (top) and ipsilateral (bottom) wing plotted against ΔF/F. Data are derived from two-minute continuous flight recordings from 20 flies. Both fluorescence and wingbeat signals have been normalized to z-scores; data are presented as mean (solid line) and a boot-strapped 95% CI for the mean (shaded area). (**F**) Correlation between GCaMP6f fluorescence and wingbeat amplitude plotted on a pixel-by-pixel basis within the recording ROI. The top image shows the expression pattern of tdTomato in the ROI of an example fly. The middle image shows the pixel-by-pixel correlation of the ΔF/F signal with left wingbeat amplitude recorded over a 2-minute flight bout; the bottom image shows the corresponding pixel-by-pixel correlation for right wingbeat amplitude. Note that activity in the DNg02 cells are positively correlated with wingbeat amplitude in the contralateral wing and negatively correlated with wingbeat amplitude of the ipsilateral wing.

## Discussion

Compared to birds, bats, and pterosaurs—the three other groups of organisms capable of sustained active flight—a unique feature of insects is that their wings are novel structures that are not modified from prior ambulatory appendages. Insects retained the six legs of their apterogote ancestors, but added two pairs of more dorsally positioned wings(16). This evolutionary quirk has profound consequences for the underlying neuroanatomy of the insect flight system. Within their thoracic ganglia, the sensory-motor neuropil associated with the wings constitutes a thin, dorsal layer sitting atop the larger ventral regions that control leg motion(6, 17, 18). Numerically, however, there appear to be more DNs targeting the flight neuropil than targeting the leg neuromeres(6). This is surprising, given the more ancient status of the leg motor system and the importance of legs in so many essential behaviors. However, the relatively large number of flight DNs may reflect the fact that control of flight requires greater motion precision because even minute changes in wing motion have large consequences on the resulting aerodynamics(19). In this paper, we describe a class of DNs in *Drosophila* (DNg02) that are unusual in that instead of existing as a unique bilateral pair, they constitute a large homomorphic population. By optogenetically driving different numbers of cells, we demonstrated that DNg02 cells can regulate wingbeat amplitude over a wide dynamic range (Figure 3A) and can elicit maximum power output from the flight motor (Figure 3D). Using 2-photon functional imaging, we also show that at least some DNg02 cells are responsive to large field visual motion during flight in a manner that would make them well suited for continuously regulating wing motion in response to both bilaterally symmetrical and bilaterally asymmetrical patterns of optic flow (Figure 4D).

Straight flight in *Drosophila* is only possible due to the maintenance of subtle and constant bilateral differences in wing motion, carefully regulated by feedback from sensory structures such as the eyes(20, 21), antennae(22, 23), and halteres(24, 25). The control system necessary for straight flight must permit the maintenance of very large, yet finely regulated, distortions of wing motion in order to produce perfectly balanced forces and moments. One means of controlling fine-scaled sensitivity over a large dynamic range is through the use of a population code with range fractionation. The use of a population code to specify motor output is a general principle(14) that has been observed in a wide array of species including leeches(26), crickets(27), cockroaches(28), and monkeys(29). In dragonflies, 8 pairs of DNs—a group of cells roughly comparable in number to the DNg02 cells—project to the flight neuropil and encode the direction to small visual targets(30).

Although the DNg02 neurons are morphologically similar, we strongly suspect that the population is not functionally homogeneous. To fly straight with perfect aerodynamic trim, an animal needs to zero its angular velocity about the yaw, pitch, and roll axes, in addition to regulating its forward flight speed, side slip, and elevation. Thus, if the DNg02 cells are the main means by which flies achieve flight trim, one would expect that they would be organized into several functional subpopulations, with each set of cells controlling a different degree of freedom of the flight motor system. For example, one subpopulation of DNg02 cells might be primarily responsible for regulating roll, while another is responsible for regulating pitch, and yet another regulates forward thrust. Such subpopulations need not constitute exclusive sets, but rather might overlap in function, collectively operating like a joystick to regulate flight pose. If this hypothesis is correct, we would expect the DNg02 neurons to differ with respect to both upstream inputs from directionally tuned visual interneurons as well as downstream outputs to power and steering muscle motor neurons. Unfortunately, we could not distinguish individual cell types across the different driver lines we used at the level of light-based microscopy. If DNg02 cells are further stratified into subclasses, it is likely that each driver line targets a different mixture of cell types. Indeed, the variation we observed in changes in wingbeat amplitude as a function of the number of DNg02 cells activated (Figure 3E) might reflect this variation in the exact complement of cells targeted by the different driver lines. Further, although one driver line (R42B02) targets 15 DNg02 neurons, it is likely that this number underestimates the size of the entire population, and we speculate that there may be a small set of neurons dedicated to regulating each output degree of freedom. Collectively, our results suggest that we have identified a critical component of the sensory motor pathway for flight control in *Drosophila,* the precise organization of which is now available for further study using a combination of genetic, physiological, and connectomic approaches.

## ACKNOWLEDGMENTS

A portion of this work was conducted as part of the Descending Interneuron Project Team at Janelia Research Campus. We would like to thank the Janelia Visiting Science Program for hosting MHD, Gudrun Ihrke and The Project Technical Resources group for assistance in coordinating the screening, and the Janelia FlyCore assisted with animal preparations. Research reported in this publication was supported by the Howard Hughes Medical Institute (S.N., W.J.R., G.M.C., W.K.) and the National Institute of Neurological Disorders and Stroke of the National Institutes of Health (I.G.R., M.H.D.) under Award U19NS104655.

## AUTHOR CONTRIBUTIONS

(Following CRediT taxonomy): Conceptualization: S.N., G.M.C., W.K, and M.H.D.; Methodology: S.N., G.M.C., W.K., and M.H.D.; Software: I.G.R., W.J.R.; Validation: W.J.R., C.M., W.K., G.M.C., I.G.R., M.H.D.; Formal Analysis: S.N., I.G.R., C.M., W.J.R., W.K., M.H.D.; Investigation: S.N., I.G.R., W.J.R.; Resources: G.M.C. W.K., M.H.D.; Data Curation: C.M., W.J.R., I.G.R.; Writing (Original Draft): S.N., M.H.D.; Writing (Review & Editing): W.K., I.G.R., M.H.D.; Visualization: S.N., I.G.R., C.M., W.K., M.H.D.; Funding Acquisition: G.M.C., W.K., M.H.D.; Supervision: G.M.C., W.K., M.H.D.; Project Administration: G.M.C., W.K., M.H.D.

## STAR METHODS

**KEY RESOURCES TABLE.**

| REAGENT or RESOURCE | SOURCE | IDENTIFIER |
| --- | --- | --- |
| **Chemicals, Peptides, and Recombinant Proteins** |  |  |
| All-trans-retinal | Sigma-Aldrich | CAS: 116-31-4 |
| Schneider's Insect Medium | Sigma-Aldrich | S0146 |
| Triton X-100 | Sigma-Aldrich | X100 |
| Xylene | Thermo Fisher Scientific | x5-500 |
| Dibutyl phthalate in xylene (DPX) | Electron Microscopy Sciences | 13512 |
| Paraformaldehyde | Electron Microscopy Sciences | 15713-S |
| **Antibodies** |  |  |
| Mouse mAb anti-Bruchpilot (nc82) | Developmental Studies Hybridoma Bank | nc82; RRID: AB_2314866 |
| rabbit polyclonal anti-GFP | Thermo Fisher Scientific | Cat #: A-11122; RRID: AB_221569 |
| Alexa Fluor 488 goat anti-rabbit | Thermo Fisher Scientific | Cat #: A-11034; RRID: AB_2576217 |
| Alexa Fluor 568 goat anti-mouse | Thermo Fisher Scientific | Cat #: A-11031; RRID: AB_144696 |
| **Deposited Data** |  |  |
| Raw and analyzed data | This paper | https://doi.org/10.17632/7q984jm2zc.1 |
| **Experimental Models: Organisms/Strains** |  |  |
| *D. melanogaster*: UAS-CsChrimson | Bloomington Drosophila Stock Center | RRID:BDSC_55135 |
| *D. melanogaster*: UAS-OpGCaMP6f (20XUAS-IVS-Syn21-OpGCaMP6F-p10 in attP5) | Gift from D. Anderson | N/A |
| *D. melanogaster*: UAS-tdTomato (P{w[+mC]=UAS-tdTom.S}3) | Bloomington | RRID:BDSC_36328 |
| *D. melanogaster*: UAS-OpGCaMP6f; UAS-tdTomato | Constructed from above two lines | N/A |
| *D. melanogaster*: pJFRC200-10XUASIVS- myr::smGFP-HA in attP18 | [(31)] | N/A |
| *D. melanogaster*: SS01074 | Bloomington Drosophila Stock Center | RRID:BDSC_75840 |
| *D. melanogaster*: SS01063 | Bloomington Drosophila Stock Center | RRID:BDSC_75837 |
| *D. melanogaster*: SS00735 | Bloomington Drosophila Stock Center | RRID:BDSC_75998 |
| D. melanogaster:<br>SS01540 | Bloomington Drosophila Stock<br>Center | RRID:BDSC_75903 |
| D. melanogaster:<br>SS01052 | Bloomington Drosophila Stock<br>Center | RRID:BDSC_75825 |
| D. melanogaster:<br>SS01558 | Bloomington Drosophila Stock<br>Center | RRID:BDSC_75946 |
| D. melanogaster:<br>SS02377 | Bloomington Drosophila Stock<br>Center | RRID:BDSC_75874 |
| D. melanogaster:<br>SS02384 | Bloomington Drosophila Stock<br>Center | RRID:BDSC_75958 |
| D. melanogaster:<br>SS01053 | Bloomington Drosophila Stock<br>Center | RRID:BDSC_86728 |
| D. melanogaster:<br>SS02631 | Bloomington Drosophila Stock<br>Center | RRID:BDSC_75976 |
| D. melanogaster:<br>SS02536 | Bloomington Drosophila Stock<br>Center | RRID:BDSC_75940 |
| D. melanogaster:<br>SS01069 | Bloomington Drosophila Stock<br>Center | RRID:BDSC_75828 |
| D. melanogaster:<br>SS02542 | Bloomington Drosophila Stock<br>Center | RRID:BDSC_75941 |
| D. melanogaster:<br>SS01546 | Bloomington Drosophila Stock<br>Center | RRID:BDSC_75944 |
| D. melanogaster:<br>SS02392 | Bloomington Drosophila Stock<br>Center | RRID:BDSC_75878 |
| D. melanogaster:<br>SS1075 | Bloomington Drosophila Stock<br>Center | RRID:BDSC_75841 |
| D. melanogaster:<br>SS01056 | Bloomington Drosophila Stock<br>Center | RRID:BDSC_75818 |
| D. melanogaster:<br>SS01556 | Bloomington Drosophila Stock<br>Center | RRID:BDSC_75953 |
| D. melanogaster:<br>SS02393 | Bloomington Drosophila Stock<br>Center | RRID:BDSC_75933 |
| D. melanogaster:<br>SS02608 | Bloomington Drosophila Stock<br>Center | RRID:BDSC_75966 |
| D. melanogaster:<br>SS01058 | [(6)] | N/A |
| D. melanogaster:<br>SS02383 | Bloomington Drosophila Stock<br>Center | RRID:BDSC_75888 |
| D. melanogaster:<br>SS02552 | Bloomington Drosophila Stock<br>Center | RRID:BDSC_75942 |
| D. melanogaster:<br>SS02379 | Bloomington Drosophila Stock Center | RRID:BDSC_75963 |
| D. melanogaster:<br>SS02635 | Bloomington Drosophila Stock Center | RRID:BDSC_75969 |
| D. melanogaster:<br>SS03500 | This paper | N/A |
| D. melanogaster:<br>SS02111 | Bloomington Drosophila Stock Center | RRID:BDSC_75935 |
| D. melanogaster:<br>SS02396 | Bloomington Drosophila Stock Center | RRID:BDSC_75882 |
| D. melanogaster:<br>SS01579 | Bloomington Drosophila Stock Center | RRID:BDSC_75902 |
| D. melanogaster:<br>SS02634 | Bloomington Drosophila Stock Center | RRID:BDSC_75970 |
| D. melanogaster:<br>SS02617 | Bloomington Drosophila Stock Center | RRID:BDSC_75967 |
| D. melanogaster:<br>SS01061 | Bloomington Drosophila Stock Center | RRID:BDSC_75836 |
| D. melanogaster:<br>SS02551 | This paper | N/A |
| D. melanogaster:<br>SS02553 | Bloomington Drosophila Stock Center | RRID:BDSC_75936 |
| D. melanogaster:<br>SS02627 | This paper | N/A |
| D. melanogaster:<br>SS01049 | Bloomington Drosophila Stock Center | RRID:BDSC_75833 |
| D. melanogaster:<br>SS01541 | Bloomington Drosophila Stock Center | RRID:BDSC_75891 |
| D. melanogaster:<br>SS01577 | This paper | N/A |
| D. melanogaster:<br>SS01578 | This paper | N/A |
| D. melanogaster:<br>SS01560 | Bloomington Drosophila Stock Center | RRID:BDSC_75948 |
| D. melanogaster:<br>SS02535 | This paper | N/A |
| D. melanogaster:<br>SS01073 | Bloomington Drosophila Stock Center | RRID:BDSC_75839 |
| D. melanogaster:<br>SS02625 | Bloomington Drosophila Stock Center | RRID:BDSC_75974 |
| D. melanogaster:<br>SS01563 | This paper | N/A |
| D. melanogaster:<br>SS02630 | This paper | N/A |
| D. melanogaster:<br>SS02550 | This paper | N/A |
| D. melanogaster:<br>SS01562 | This paper | N/A |
| D. melanogaster:<br>GMR42B02 | Bloomington Drosophila Stock<br>Center | RRID:BDSC_68782 |
| D. melanogaster:<br>SS02624 | Bloomington Drosophila Stock<br>Center | RRID:BDSC_75972 |
| D. melanogaster:<br>SS02544 | This paper | N/A |
| **Software and Algorithms** |  |  |
| Kinefly wing tracking software | [(9)] | https://github.com/ssafarik/Kinefly |
| Python | https://www.python.org | RRID: SCR_008394 |
| Matplotlib | https://matplotlib.org | RRID: SCR_008624 |
| MATLAB | https://www.mathworks.com | RRID: SCR_001622 |
| FIJI | https://fiji.sc/ | RRID:SCR_002285 |
| **Other** |  |  |
| UV-activated cement | Henkel | Loctite® 3972TM |
| Digital camera | Balser | acA640–120 gm |
| Long-pass filter | Midwest Optical Systems | LP715-30.5 |
| Fiber optic light guide | Thorlabs | FT1500EMT |
| IR light source LED | Thorlabs | M850F2 |
| IR light source driver | Thorlabs | LEDD1B |
| Wingbeat analyzer | Phidgets | PhidgetAnalog 1002 |
| Data acquisition system | Axon Instruments | Digitata 1440A |
| Panel controller | IO Rodeo | Panels display controller unit |
| Transmission filter | Indigo | Roscolux no. 59 filter |
| Transmission filter | Skelton Exotic Sangria | Rosco no. 39 filter |
| Transmission filter | Cyan | Rosco #4390 filter |

## RESOURCE AVAILABILITY

### Lead contact

Further information and requests for resources and reagents should be directed to and will be fulfilled by the Lead Contact, Michael H. Dickinson.

### Materials availability

All new fly lines generated for this paper are listed in the key resources table.

### Data and code availability

All data has been deposited on Mendeley at https://doi.org/10.17632/7g984jm2zc.1 and are publicly available as of the date of publication. The DOI is listed in the key resources table. All original code for data analysis has been deposited on Mendeley at https://doi.org/10.17632/7g984jm2zc.1 and is publicly available as of the date of publication. All original code used for the kinefly wing tracking software is publicly available on Github at https://github.com/ssafarik/Kinefly. The DOIs are listed in the key resources table. Any additional information required to reanalyze the data reported in this paper is available from the lead contact upon request.

## EXPERIMENTAL MODEL AND SUBJECT DETAILS

All experiments were conducted on genetically modified female *Drosophila melanogaster.* To create genetic lines that specifically targeted DNg02 cells, we used the split-GAL4 technique described previously (Luan et al. 2006; Namiki et al. 2018). We combined split half lines that have promoters to drive expression of either the transcription activation domain (p65ADZp) or the DNA-binding domain (ZpGAL4DBD) of the GAL4 protein. To identify driver lines containing DNg02, we manually searched the brain expression pattern from publically available GAL4 lines in the FlyLight database (https://flweb.janelia.org/). We chose pairs of these driver lines that appeared to target the DNg02 cells and screened the resulting split-GAL4 combinations by crossing them with flies carrying the reporter pJFRC200-10XUASIVS-myr::smGFP-HA inserted into attP18. For the optogenetic activation experiments, we used 3-to-6-day-old female flies obtained by crossing virgin females from each split-GAL4 line (or GMR42B02) with 3-to-5-day-old males carrying 20XUAS-CsChrimson-mVenus inserted into attP18. We reared the progeny of this cross on standard cornmeal fly food containing 0.2 mM all trans-Retinal (ATR) (Sigma-Aldrich) and transferred adult flies 0-2 days after eclosion onto standard cornmeal fly food with 0.4 mM ATR.

We supplemented the standard cornmeal food with additional yeast. For functional imaging experiments, we used 2-5 days old female flies that resulted from a cross of the split-GAL4 line SS02535 with w+;UAS-tdTomato;UAS-GCaMP6f flies(32).

## QUANTIFICATION AND STATISTICAL ANALYSIS

All experiments were analyzed with custom written scripts written in either Matlab or Python. This manuscript includes no explicit tests of significance. Sample sizes refer to the number of individuals tested.

## METHODS DETAILS

### Optogenetic activation experiments

Female offspring of split-GAL4 driver lines crossed to flies carrying 20XUAS-CsChrimson-mVenus were anesthetized on a cold surface (4°C) and tethered to a tungsten pin with Loctite^®^ 3972™ UV-activated cement (Henkel). The tethered fly was positioned such that its stroke plane was horizontal and perpendicular to the vertical optical axis of a digital camera (Basler acA640– 120 gm) equipped with an Infinistix 90-degree lens with a long-pass filter (LP715-30.5, Midwest Optical Systems). Two horizontally oriented fiber optic light guides (FT1500EMT; Thorlabs), each coupled to an IR light source (driver: LEDD1B, and LED: M850F2; Thorlabs) illuminated the stroke planes of the left and right wings. We used Kinefly software(9) to track the anterior-most angular excursion of the fly’s left and right wingbeat (Figure 1C). Digital values for the left and right wingbeat amplitudes were converted into voltages using a PhidgetAnalog 1002 (Phidgets), and recorded on a Digitata 1440A data acquisition system (Axon Instruments) for subsequent analysis. The voltage signals from the two wings were also sent to an LED panel controller(10) (IOrodeo) which was programmed so that flies could regulate the angular velocity of the visual display via the difference in wingbeat amplitude of the two wings. We used a custom-built photodetector circuit to record the oscillations in the incident IR light caused by the flapping motion of the wings and convert that signal into a voltage proportional to wingbeat frequency (https://github.com/janelia-kicad/light_sensor_boards), which we also recorded on the Digidata 1440A. For optogenetic activation, we positioned a fiber optic light guide (FT1500EMT; Thorlabs) beneath the fly, aimed at the thorax, which conducted the output of a 617 nm LED (M617F1, Thorlabs) at ~3.4 mW/mm^2^. The timing and duration of the pulse was controlled via the voltage input to the LED driver (LEDD1B, Thorlabs). The fly and ancillary instruments were surrounded by a 12 × 4 panel (96 × 32 pixel) LED arena (470 nm)(10) that covered 216° of azimuth with a resolution of 2.25° in front of the fly. In each experiment, we elicited 30 responses to a 100 ms light pulse with an interpulse interval of 10 seconds under visual closed loop conditions, in which the difference in fly’s left-right wingbeat amplitude controlled the angular velocity of a striped drum with a spatial frequency of 36°. For each trial, we determined the average wingbeat amplitude (WBA) of the left and right wings over the 0.5 second period prior to stimulus onset and subtracted this value from the entire trace to create a zero baseline. The response to optogenetic activation was calculated as the average value of WBA over the 0.5 second period starting with stimulus onset. The 30 trails from each fly were averaged to create a single measurement for each individual.

### Anatomy

To image expression patterns, we dissected the complete central nervous systems of 3-to-5-day-old female adult progeny in Schneider’s Insect Medium (Sigma), fixed them in paraformaldehyde, and then transferred them to a goat serum blocking buffer for 1 hr. We then replaced the buffer with the primary antibodies (mouse nc82 supernatant at 1:30, rabbit polyclonal anti-GFP at 1:1000) diluted in phosphate buffered saline with 0.5% Triton X-100 (PBT) and gently agitated the preparations for 36–48 hr at 4°C. After washing with PBT, the samples were then incubated with secondary antibodies (Alexa Fluor 488 goat anti-rabbit, and Alexa Fluor 568 goat anti-mouse at 1:400) diluted in PBT and agitated again at 4°C for 3 days. The samples were then washed, fixed again in paraformaldehyde, mounted on a poly-L-lysine cover slip, cleared with xylene, and embedded in dibutyl phthalate in xylene (DPX) on a microscope slide with spacers. After drying for two days, samples were imaged at either 20X or 40X with a confocal microscope (Zeiss LSM 510) (Dionne et al. 2018).To discriminate the morphology of individual DNg02 cells, we used a multi-color flip out technique(12). To identify pre-synaptic terminal of DNg02, we examined neuronal polarity using a reporter (pJFRC51-3xUAS-Syt::smGFP-HA in su(Hw)attPa) that labels synaptotagmin, a synaptic vesicle-specific protein. More detailed descriptions of these protocols are available on the Janelia FlyLight website (https://www.janelia.org/project-team/flylight/protocols). All images are available through the FlyLight Split-GAL4 website (https://splitgal4.janelia.org/cgi-bin/splitgal4.cgi).

### Functional Imaging

Tethered, flying flies were imaged at an excitation wavelength of 930 nm using a galvanometric scan mirror-based two-photon microscope (Thorlabs) equipped with a Nikon CFI apochromatic, near-infrared objective water-immersion lens (40x mag., 0.8 N.A., 3.5 mm W.D.). We used the 2-5 days old female progeny of a cross between the split-GAL4 driver line SS02535, which drives expression in 3 pairs of DNg02s and w+;UAS-tdTomato;UAS-GCaMP6f flies(32). We recorded two channels to image tdTomato and GCaMP6f fluorescence in the posterior slope arbors of both left and right DNg02 neurons. Depending on the individual preparation, we acquired either 72 × 36 μm images with 128 × 64 pixel resolution at 13.1 Hz, or 72 × 29 μm images with 160 × 64 pixel resolution at 11.2 Hz. To correct for horizontal motion in the x-y plane, we registered both channels for each frame by finding the peak of the cross correlation between each tdTomato image and the trial-averaged image(33). Based on tdTomato expression we selected a field of view (FOV) approximately centered along the medio-lateral axis. The 20% most variable pixels in the left and right halves of the GCaMP6f images were selected as the left and right regions of interest (ROIs) respectively. To correct for motion in z, we normalized the GCaMP6f fluorescence to the tdTomato fluorescence. For each frame and each side, we computed fluorescence (F_t_) of the GCaMP6f signal by subtracting the average of the background from the average of the ROI. The background was defined as the 20% dimmest pixels of the entire FOV. We computed the baseline fluorescence, F_0_, as the mean of the 10% lowest F_t_ in the ROI. To standardize the measured neuronal activity across individual preparations we normalized baseline-subtracted fluorescence to the maximum observed for each individual fly on each anatomical side of the brain as ΔF/F = (F_t_ – F_0_) / (F_t_ – F_0_) _max_. We presented visual stimuli with a 12 × 4 panel (96 × 32 pixel) arena that covered 216° of azimuth with a resolution of 2.25°, identical to that used in the optogenetic screen. To reduce light pollution from the LED arena into the photomultiplier tubes of the 2-photon microscope, we shifted the spectral peak of the visual stimuli from 470 nm to 450 nm by placing five transmission filters in front of the LEDs (one sheet of Roscolux no. 59 Indigo, two sheets of no. 39 Skelton Exotic Sangria, and two sheets of no. 4390 Cyan). We presented an array of visual patterns, each for 3 s, alternated with 3 s static starfields. The visual patterns were presented in a shuffled pseudo-random order and included roll, pitch, and yaw motion in both directions, a stripe oscillating on the left or right, an expanding object on the left or right, progressive and regressive motion, and closed loop stripe fixation. We illuminated the wings using four horizontal fiber-optic IR light sources (M850F2, Thorlabs) distributed in a ~90° arc behind the fly. We tracked left and right wingbeat amplitudes with a machine vision system, Kinefly(9), at 32 Hz. This method introduces a delay in measurement of ~30 ms, which we corrected.

